# Nitrate reduction stimulates and is stimulated by phenazine-1-carboxylic acid oxidation by *Citrobacter portucalensis* MBL

**DOI:** 10.1101/2021.06.04.447179

**Authors:** Lev M. Tsypin, Dianne K. Newman

**Affiliations:** Division of Biology and Biological Engineering, California Institute of Technology, Pasadena, CA, USA; Division of Geological and Planetary Sciences, California Institute of Technology, Pasadena, CA, USA

## Abstract

Phenazines are secreted metabolites that microbes use in diverse ways, from quorum sensing to antimicrobial warfare to energy conservation. Phenazines are able to contribute to these activities due to their redox activity. The physiological consequences of cellular phenazine reduction have been extensively studied, but the counterpart phenazine oxidation has been largely overlooked. Phenazine-1-carboxylic acid (PCA) is common in the environment and readily reduced by its producers. Here, we describe its anaerobic oxidation by *Citrobacter portucalensis* strain MBL, which was isolated from topsoil in Falmouth, MA, and which does not produce phenazines itself. This activity depends on the availability of a suitable terminal electron acceptor, specifically nitrate. When *C. portucalensis* MBL is provided reduced PCA and nitrate, it rapidly oxidizes the PCA. We compared this terminal electron acceptor-dependent PCA-oxidizing activity of *C. portucalensis* MBL to that of several other γ-proteobacteria with varying capacities to respire nitrate. We found that PCA oxidation by these strains in a nitrate-dependent manner is decoupled from growth and correlated with their possession of the periplasmic nitrate reductase Nap. We infer that bacterial PCA oxidation is widespread and propose that it may be genetically determined. Notably, oxidizing PCA enhances the rate of nitrate reduction to nitrite by *C. portucalensis* MBL beyond the stoichiometric exchange of electrons from PCA to nitrate, which we attribute to *C. portucalensis* MBL’s ability to also reduce oxidized PCA, thereby catalyzing a complete PCA redox cycle. This bidirectionality highlights the versatility of PCA as a biological redox agent.

**IMPORTANCE:** Phenazines are increasingly appreciated for their roles in structuring microbial communities. These tricyclic aromatic molecules have been found to regulate gene expression, be toxic, promote antibiotic tolerance, and promote survival under oxygen starvation. In all of these contexts, however, phenazines are studied as electron acceptors. Even if their utility arises primarily from being readily reduced, they need to be oxidized in order to be recycled. While oxygen and ferric iron can oxidize phenazines abiotically, biotic oxidation of phenazines has not been studied previously. We observed bacteria that readily oxidize phenazine-1-carboxylic acid (PCA) in a nitrate-dependent fashion, concomitantly increasing the rate of nitrate reduction to nitrite. Because nitrate is a prevalent terminal electron acceptor in diverse anoxic environments, including soils, and phenazine-producers are widespread, this observation of linked phenazine and nitrogen redox cycling suggests an underappreciated role for redox-active secreted metabolites in the environment.

## OBSERVATION

Physiological studies of phenazines have focused on cellular reduction of these secreted molecules for over 120 years. Reduction of phenazines by bacteria was first proposed in the 19^th^ century as an indicator for the presence of enteric bacteria in water supplies (1). Several decades later, pyocyanin, one of the phenazines produced by *Pseudomonas aeruginosa*, was described as an “accessory respiratory pigment” that increased the rate of oxygen consumption by *Staphylococcus*, *Pneumococcus*, and erythrocytes by shuttling electrons from the cells to oxygen (2). Once it became apparent that phenazines can have cytotoxic effects, they were characterized as antimicrobial compounds that destructively abstract electrons from the transport chain (3). It was then discovered that phenazine reduction can greatly benefit *P. aeruginosa* by: 1) regulating gene expression in *P. aeruginosa* during quorum sensing by oxidizing a transcription factor; 2) acting as alternative terminal electron acceptors to promote anoxic survival; and 3) facilitating iron acquisition (4–8). These reports paint a complex picture of the multifarious effects phenazines can have, but in each case the conceptual model ends with the cell reducing the phenazine. Reduced phenazines can be re-oxidized by inorganic electron acceptors like oxygen and ferric iron, and this abiotic process has been invoked to explain redox cycling of phenazines in biofilms (9, 10). However, when these electron acceptors are unavailable, biotic oxidation of reduced phenazines could close the redox cycle by regenerating oxidized phenazines. This process has not been shown to exist for secreted redox-active metabolites.

In parallel to these physiological studies, phenazines have been used as generic electron shuttles in bioelectrochemical reactor research (BER), selected according to their chemical properties and suitability for a given application (11, 12). Electrochemically reduced neutral red (NR), a phenazine, has been successfully used as an electron donor to cells, chosen for its standard midpoint potential (very near to that of NADH/NAD^+^, 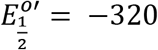 mV vs. NHE) and hydrophobicity (13–15). Anaerobic NR oxidation in BERs has been coupled to the reduction of several terminal electron acceptors, including nitrate (16). A limitation of these studies is that NR is not found in nature. Therefore, despite NR oxidation being useful in regulating electrosynthesis, the existence of natural bacterially driven phenazine oxidation remains unexplored.

PCA is one of the mostly widely synthesized phenazines in the microbial world, from which other phenazines are derived (17, 18). PCA is known to be reduced by its producers, driving current generation in bioelectrochemical systems, in which it is re-oxidized by the anode (5, 19). These facts make PCA a fitting candidate for microbial oxidation during anaerobic metabolism.

In previous work, we enriched for PCA oxidizers from topsoil by incubating them with reduced PCA, acetate (a non-fermentable carbon source), and nitrate as the only terminal electron acceptor, and successfully isolated the PCA-oxidizing *Citrobacter portucalensis* MBL, which unable to synthesize its own phenazines (20).

In this Observation, we performed three sets of experiments to study biological redox transformations of PCA: an oxidation assay in which reduced PCA (PCA_red_) was incubated with or without cells and with or without a terminal electron acceptor (Fig. 1); a reduction assay in which oxidized PCA (PCA_ox_) was incubated with or without cells and with or without a terminal electron acceptor (Supp. Fig. 1); and an ion chromatography experiment in which we measured the reduction of nitrate to nitrate by *C. portucalensis* MBL depending on PCA_red/ox_ availability (Fig. 2A).

**Figure 1.**
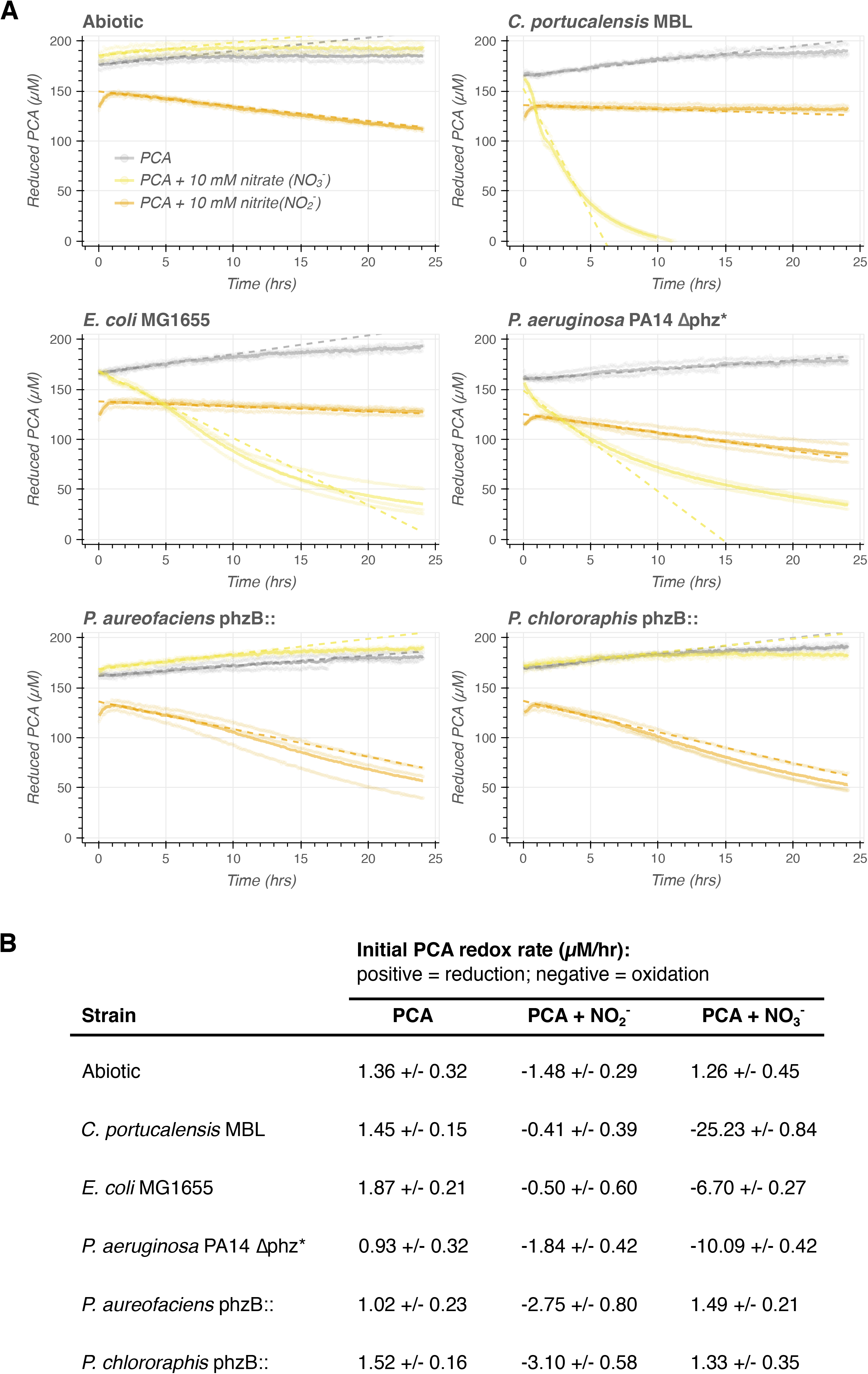
Oxidation of PCA_red_ by bacteria provided different terminal electron acceptors (TEAs). (A) The light circles correspond to three independent biological replicates and the dark lines to their respective means. Cells only oxidize PCA when an appropriate TEA is available. Nitrate and nitrite (yellow and orange curves, respectively) stimulate different strains to oxidize PCA_red_. When no TEA is provided (grey curves), no strains oxidize PCA_red_. With nitrate, only *P. chlororaphis* and *aureofaciens* appear to not oxidize PCA_red_. Nitrite abiotically oxidizes PCA_red_ (orange curve, top left panel), but *P. aureofaciens and chlororaphis* catalyze an even faster biological oxidation. In contrast, the *Enterics* (*C. portucalensis* MBL and *E. coli* MG1655) reduce PCA_ox_ faster than the abiotic reaction with nitrite can compensate. The dashed lines correspond to the linear fits reported in (B). (B) This table reports the estimated initial rates of oxidation according to a linear fit over the first five hours. This timeframe was determined by tracking the R^2^ for the linear fit over increasing time windows (Supp. Fig. 4). PCA_ox_ reduction is calculated as a positive rate; PCA_red_ oxidation is calculated as a negative rate.

**Figure 2.**
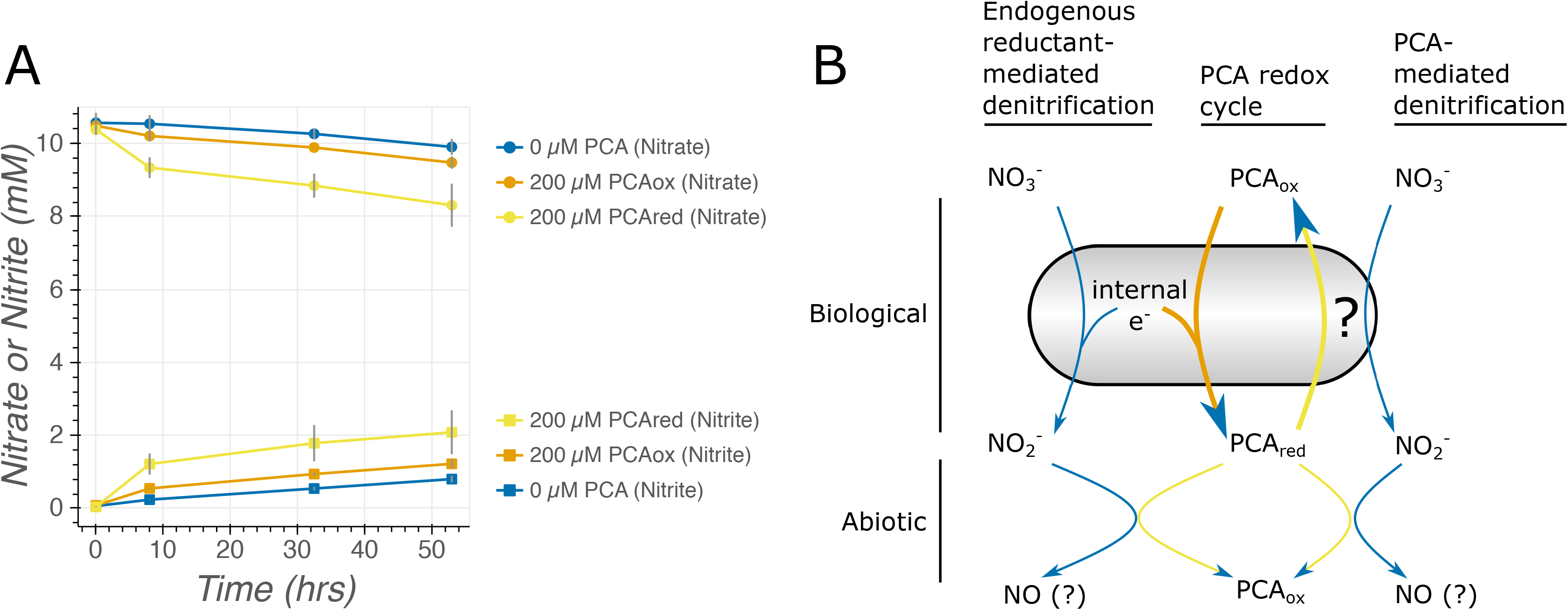
PCA oxidation by *C. portucalensis* MBL increases its initial rate of nitrate reduction. (A) Either 200 μM reduced PCA (PCA_red_), 200 μM oxidized PCA (PCA_ox_), or no PCA was added to each condition. Ion chromatography shows that over the 10 hours that the *C. portucalensis* MBL cells are oxidizing PCA_red_ (Fig. 1A), their initial rate of nitrate reduction is substantially increased. The nitrate is stoichiometrically reduced to nitrite. Error bars are 95% confidence intervals around the mean of three independent biological replicates. When not visible, the intervals are smaller than the circles (nitrate) or squares (nitrite) denoting the measurements. (B) Blue arrows denote denitrification, yellow arrows denote observed paths of PCA_red_ oxidation, and the orange arrow denotes the observed path of PCA_ox_ reduction. Any cell that has internal stores of reducing equivalents and an appropriate terminal electron acceptor (for *C. portucalensis* MBL, *E. coli* MG1655, and *P. aeruginosa* Δphz* in our experiments—nitrate) may catalyze an internal PCA redox cycle (bolded arrows). *P. aureofaciens* and *chlororaphis* may also do this with nitrite. In addition to this cellularly-catalyzed reaction, the product of nitrate’s reduction (nitrite) may abiotically oxidize PCA (Fig. 1A, top left panel, orange curve). The nature of the biological oxidation of PCA coupled to nitrate reduction remains unknown but will be amenable to genetic experiments.

### *C. portucalensis* MBL oxidizes PCA in a nitrate-dependent manner

We did not observe PCA (PCA_red_/PCA_ox_ 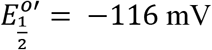 vs. NHE (9)) oxidation in the absence of a terminal electron acceptor in either the abiotic and biotic regimes (Fig. 1A, grey curves; Fig. 1B, PCA-only column). When 10 mM nitrate (NO_3_^−^/NO_2_^−^ 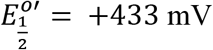 vs NHE (15)) was added (right panels), *C. portucalensis* MBL readily oxidized PCA_red_ at an initial rate of 25.23 ± 0.84 μM/hr (Fig. 1A, top right panel, yellow curve; Fig. 1B). Nitrate did not oxidize PCA abiotically (Fig. 1A, top left panel, yellow curve). During both dissimilatory and assimilatory nitrate reduction, nitrate is first reduced to nitrite (21). Nitrite (NO_2_^−^/NO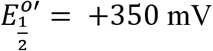 vs NHE (15)) drove slow abiotic oxidation of PCA_red_ at a steady rate of −1.48 ± 0.29 μM/hr after an initial short period of reduction (Fig. 1A, top left panel, orange curve; Fig. 1B). This abiotic rate of oxidation is insufficient to be responsible for the rate of PCA_red_ oxidation by cells in the presence of nitrate.

When *C. portucalensis* MBL is incubated with PCA_red_ and nitrite, there is less PCA_red_ oxidation than in the abiotic case (Fig. 1A, top right panel, orange curve). We interpret this to mean that *C. portucalensis* catalyzes the oxidation of PCA_red_ when an appropriate terminal electron acceptor is available but reduces PCA_ox_ when such an electron acceptor is absent. We observed PCA_ox_ reduction when the cells started with 200 μM PCA_ox_, but the rate of reduction decreased according to the provided terminal electron acceptor (Supp. Fig. 1). Adding nitrate to the PCA_ox_ condition caused reduction to be non-detectable; and the presence of nitrite, which abiotically oxidizes PCA_red_, caused a slight dampening in the reduction rate by 1.25 ± 0.09 μM/hr (Supp. Fig. 1).

### Comparative study of nitrate- and fumarate-dependent PCA oxidation by several γ-proteobacteria

We assayed whether other γ-proteobacteria can also oxidize PCA in a terminal electron acceptor-dependent manner (Fig. 1). Using the same assay as above, *E. coli MG1655*, *P. aeruginosa* UCBPP-PA14 *Δphz\**, *P. chlororaphis phzB::TnluxAB*, and *P. aureofaciens phzB::lacZ* (all of which cannot synthesize PCA either naturally or due to the specified mutations) were incubated with PCA and 10 mM nitrate or nitrite, or no terminal electron acceptor. With the exception of *P. chlororaphis* and *aureofaciens*, all strains exhibited PCA oxidation with nitrate, with *C. portucalensis* MBL being the fastest (Fig. 1B). The assayed *P. aureofaciens* and *chlororaphis* oxidized PCA_red_ with nitrite faster than the abiotic control: −2.75 ± 0.80 and −3.10 ± 0.58, respectively, vs. −1.48 ± 0.29 μM PCA_red_/hr abiotically (Fig. 1B). The reduction assay for the other strains reflected our observations with *C. portucalensis* MBL: when a terminal electron acceptor that stimulated PCA_red_ oxidation was present, no reduction was observed (Supp. Fig. 1). None of the strains exhibited significant growth in these assays (Supp. Figs. 2 and 3).

### The effect of PCA oxidation on the initial rate of nitrate reduction by *C. portucalensis* MBL

In Fig. 1A it is clear that *C. portucalensis* MBL completes its oxidation of ~200 μM PCA_red_ within 10 hours when nitrate is available. We repeated this experiment in anaerobic culture tubes inoculated with *C. portucalensis* MBL, 10 mM NO_3_^−^, and either 200 μM PCA_red_, 200 μM PCA_ox_, or no PCA and measured nitrate and nitrite concentrations over time via ion chromatography. We observed that PCA_red_ oxidation significantly increased the rate of nitrate reduction to nitrite (Fig. 2A). Nitrate consumption was stoichiometrically matched by nitrite production. We did not observe the production of any other nitrogen oxides or ammonium (data not shown). The no-PCA control did not show any nitrate reduction over the first eight hours (3 ± 55 μM/hr; 95% confidence interval reported for all rate measurements, calculated from a linear regression of the data from the first two timepoints for each curve in Fig. 2A). During this time, 131 ± 49 μM/hr nitrate was reduced in the PCA_red_ condition. In contrast, the PCA_ox_ control exhibited only a putative increase in nitrate reduction (35 ± 35 μM/hr). To verify that nitrate was reduced to nitrite, we tracked nitrite’s production. We observed that over the first eight hours the no-PCA control produced nitrite at the rate of 22 ± 3 μM/hr, the PCA_ox_ control at 58 ± 2 μM/hr, and the PCA_red_ condition at 147 ± 44 μM/hr of nitrite over the first eight hours. Thus, we estimate the effect of PCA_red_ vs PCA_ox_ to be 96 ± 60 μM/hr of increased nitrate reduction or 89 ± 44 μM/hr of increased nitrite production. The increase in nitrate reduction due to PCA_red_ was greater than the absolute number of electrons the PCA_red_ could provide: PCA redox and nitrate reduction are both two-electron processes (9, 21), and a process without a redox cycle would predict that oxidizing PCA_red_ at a rate of 25 μM/hr (Fig. 1B) would reduce at most 25 μM/hr nitrate to nitrite. However, the lowest range of the confidence intervals suggests that at least an extra 36 μM/hr nitrate was reduced by the cells when PCA_red_ was provided, implying that the PCA is stimulating nitrate reduction by some other means than a one-to-one electron donation; our data showing PCA reduction capability (Supp. Fig. 1) suggests that the cells may be cycling PCA under these conditions, though this conclusion awaits definitive demonstration.

### Conclusions

The effect of PCA oxidation by *C. portucalensis* MBL on its rate of nitrate reduction was outsized (Fig. 2A). This is consistent with two explanations: 1) a prior report argues that neutral red (a synthetic phenazine) oxidation affects electrosynthesis during anaerobic respiration primarily by changing gene regulation via menaquinone reduction (16), and so it is plausible that PCA oxidation may increase transcription of a rate-limiting factor in the electron transport chain to nitrate; 2) we observed that a PCA redox cycle by *C. portucalensis* MBL is possible: the cells naturally reduce PCA_ox_, but this is not detectable while nitrate is present, potentially meaning that the PCA is being re-oxidized as soon as it is reduced. If this is the case, it is possible that PCA redox cycling may also shunt more electrons to nitrate reduction. An alternative explanation for why PCA_ox_ reduction is not observed in the presence of nitrate is that the cells are preferentially reducing nitrate instead of PCA. While we cannot directly test this hypothesis given our current experimental set up, this will be possible once we develop methods to genetically modify *C. portucalensis* MBL. We have not been able to determine whether reduced PCA (PCA_red_) serves as an effective electron donor to the cells’ metabolism from our above observations. Even if there is no direct physiological benefit from PCA oxidation, that cells can catalyze this reaction and stimulate nitrate reduction has environmental ramifications far beyond the cell.

Despite all being capable of nitrate reduction, not all the γ-proteobacteria that we tested were capable of oxidizing PCA_red_ with nitrate (Fig. 1), suggesting there may be a genetic basis for this process, specifically, possession of the periplasmic nitrate reductase, NAP. The only strains that did not readily oxidize PCA_red_ with nitrate were *P. chlororaphis* and *aureofaciens*, and, unlike the other species, they do not possess the periplasmic nitrate reductase operon *nap*. Curiously, it has been previously noted in a PhD thesis that wildtype *P. aeruginosa* UCBPP-PA14 oxidizes another phenazine, pyocyanin, in the presence of nitrate, while a *napA* transposon knockout strain does not (22). We propose that this operon may be necessary for rapid PCA_red_ oxidation. An alternative possibility is that there are other membrane-bound or secreted oxidants that some of the species can produce, or that our experimental conditions did not stimulate *P. aureofaciens* or *P. chlororaphis* to exhibit the activity, despite having the capacity to do so. We are developing genetic tools in *C. portucalensis* MBL to test these hypotheses directly.

Nitrate-dependent PCA_red_ oxidation is likely to be common in anoxic environments, such as soils, as it is not species-specific and occurs readily. If the outsized effect of PCA_red_ oxidation on nitrate reduction that we observe generalizes to other species, the production and reduction of phenazines by organisms like *Pseudomonads* likely affects the rate of nitrate consumption in their environs, adding another function to the phenazine arsenal (18). We propose that cells may catalyze a PCA redox cycle whenever they have internal stores of reducing equivalents and a usable terminal electron acceptor (Fig 2B). While both intracellular (e.g., sulfur) and extracellular (e.g., humics) bacterial redox cycles have been described (23, 24), to our knowledge this has not been appreciated for secreted redox active metabolites, such as phenazines. Our observation underscores the possibility that these molecules may act as “electron buffers”, enabling cells to reduce and oxidize them according to whether they are lacking a terminal electron acceptor or an electron donor, respectively, and in so doing significantly impact other biogeochemical cycles, such as the nitrate cycle.

## MATERIALS AND METHODS

### Strains and media

*Citrobacter portucalensis* MBL was isolated in our previous work (20). In the comparative PCA oxidation and reduction experiments, we used strains of γ-proteobacteria that cannot synthesize phenazines, either natively or due to mutations. We used *E. coli* MG1655, *P. aeruginosa* UCBPP-PA14 *Δphz\** (25), *P. chlororaphis phzB::TnLuxAB* (strain PCL1119 obtained from G. Bloemberg, Leiden University (26)), and *P. aureofaciens phzB::lacZ* (strain 30-84Z obtained from L. Pierson, University of Arizona (27)). The wildtype *Pseudomonads* can synthesize PCA, but the used mutants cannot. All strains were grown and incubated under the same conditions. The basal medium for the experiments contained 20 mM potassium phosphate buffer (final pH 7), 1 mM sodium sulfate, 10 mM ammonium chloride, 1x SL-10 trace elements, 1x freshwater salt solution (17.1 mM sodium chloride, 1.97 mM magnesium chloride, 0.68 mM calcium chloride, and 6.71 mM potassium chloride), and 1x 13-vitamin solution (10 μM MOPS pH 7.2, 0.1 μg/mL riboflavin, 0.03 μg/mL biotin, 0.1 μg/mL thiamine HCl, 0.1 μg/mL L-ascorbic acid, 0.1 μg/mL d-Ca-pantothenate, 0.1 μg/mL folic acid, 0.1 μg/mL nicotinic acid, 0.1 μg/mL 4-aminobenzoic acid, 0.1 μg/mL pyridoxine HCl, 0.1 μg/mL lipoic acid, 0.1 μg/mL NAD^+^, 0.1 μg/mL thiamine pyrophosphate, and 0.01 μg/mL cyanocobalamin). Depending on the experimental condition, as indicated in the figure legends, a terminal electron acceptor would be added (10 mM of fumarate, nitrate, or nitrite) or omitted. For oxidized PCA, a 10 mM stock in 20 mM NaOH was prepared. For reduced PCA, an 800 μM stock in the basal medium was reduced by electrolysis. Both stocks were diluted into plates to a final target concentration of 200 μM.

### Cell preparation

All cell incubations and experiments were performed at 30 C. Cells were preserved in 35% glycerol stocks at −80 C. Two days prior to the experiments, frozen cells for each strain assayed were struck out on lysogeny broth (LB) agar plates and incubated overnight. The evening prior to the experiment, a patch from the streaks was inoculated into liquid LB in a respective culture tube and incubated slanted, shaking at 250 rpm, overnight. The morning of the experiment, 1 mL of each cell culture was washed three times into the basal medium by spinning for two minutes at 6000 x g, aspirating the supernatant, and gently resuspending with a pipette. The OD_600_ of each washed culture was measured. The cultures were brought into a Coy glove box, where they were washed three times into the same basal medium that had been made anoxic, following the same procedure as above. After being left to stand for 1-2 hours, the cells were inoculated into the different experimental conditions at a target starting OD_600_ of 0.1.

### Measurement of PCA redox and nitrogen oxide concentrations

All PCA redox measurements were performed in a Coy chamber (5% hydrogen/95% nitrogen headspace) using a BioTek Synergy 5 plate reader. Reduced PCA concentration was measured by fluorescence (excitation 360 nm and emission 528 nm) (28). In these cultures, 1 mM acetate was provided. Plates were incubated shaking on the “medium” setting. For nitrogen oxide concentration measurements, *C. portucalensis* MBL cells were prepared as above, but incubated in culture tubes in the Coy chamber to allow for sampling. In these cultures, no acetate was provided. The tubes were kept at 30 C, but not shaking. Samples were filtered through a 0.2 μm cellulose-acetate spin filter and stored at −80 C prior to analysis. Nitrate and nitrite concentrations were measured by ion chromatography using a Dionex ICS-2000 instrument.

### Data analysis

Initial redox rates in the conditions with supplied nitrite did not include the first 1.5 hours of the assay because that period showed a systematic artifact due to fluorescence quenching (Fig. 1A, lower left panel). The linear fits were calculated over the first five hours after the first detectable PCA_red_ measurement, which was determined to be appropriate based on scanning for R^2^ values over increasing time frames (Supp. Figs. 4 and 5). 95% confidence intervals are calculated as the estimated value ± 1.96 * (standard error). When comparing rates, the reported error is the geometric mean of the intervals for the two measurements. All plots were generated using Bokeh and the legends and titles were adjusted using Inkscape. All the raw data and the Jupyter notebook used for their analysis are available at https://github.com/ltsypin/Cportucalensis_observation.

## ACKNOWLEDGEMENTS

We would like to thank the members of the Newman lab, and especially Scott Saunders, Darcy McRose, Avi Flamholz, John Ciemniecki, Chelsey VanDrisse, and Justin Bois for their insight and helpful discussions throughout this work. We are grateful to Nathan Dalleska at the Water and Environment Laboratory at Caltech for training LMT on the Dionex instrument and providing a facility for analytical chemistry. LMT was supported by the Rosen Endowment Fellowship at Caltech and the National Science Foundation Graduate Research Fellowship (DGE-1745301). Additional support to DKN came from NIH (1R01AI127850-01A1 and 1R01HL152190-01) and ARO (W911NF-17-1-0024) grants.

## Supplementary Figure Legends

**Supplementary Figure 1. Reduction of PCA_ox_ by bacteria provided different terminal electron acceptors (TEAs).** (A) The light circles correspond to three independent biological replicates and the dark lines to their respective means. Cells only appear to reduce PCA_ox_ when there is no PCA_red_-oxidation stimulating TEA present. Nitrate (yellow curves) inhibits *C. portucalensis* MBL, *E. coli* MG1655, and *P. aeruginosa* Δphz* from reducing PCA_ox_. Nitrite inhibits PCA_ox_ reduction by the three *Pseudomonads*. When no TEA is provided (grey curves), all strains reduce PCA_ox_. There is no detectable abiotic reduction. The dashed lines correspond to the linear fits reported in (B). (B) This table reports the estimated initial rates of oxidation according to a linear fit over the first five hours. This timeframe was determined by tracking the R^2^ for the linear fit over increasing time windows (Supp. Fig. 5). PCA reduction is calculated as a positive rate; PCA oxidation is calculated as a negative rate.

**Supplementary Figure 2.** OD_600_ measurements for all strains in all conditions during the oxidation assays (Fig. 1). Light circles represent all measurements from three independent biological replicates corresponding to the PCA_red_ measurements in Figure 1. The solid lines are the means of the replicates. The curves are colored by condition (i.e., which, if any, terminal electron acceptor was added). While there are some OD_600_ trends in the first five hours, there is no evidence of growth that would indicate a doubling. In the abiotic graph, all the data points overlie each other.

**Supplementary Figure 3.** OD_600_ measurements for all strains in all conditions during the reduction assays (Supp. Fig. 1). Light circles represent all measurements from three independent biological replicates corresponding to the PCA_red_ measurements in Supp. Fig. 1. The solid lines are the means of the replicates. The curves are colored by condition (i.e., which, if any, terminal electron acceptor was added). While there are some OD_600_ trends in the first five hours, there is no evidence of growth that would indicate a doubling. In the abiotic graph, all the data points overlie each other.

**Supplementary Figure 4.** Evaluation of the appropriate window for performing the linear regressions to estimate initial redox rates in the oxidation assays (Fig. 1B). Starting from the first valid PCA_red_ measurement in each condition, a linear regression was performed over increasing lengths of time and the corresponding R^2^ value was plotted. A time window of five hours (red line) was selected as a systematic compromise for all conditions.

**Supplementary Figure 5.** Evaluation of the appropriate window for performing the linear regressions to estimate initial redox rates in the reduction assays (Supp. Fig. 1B). Starting from the first valid PCA_red_ measurement in each condition, a linear regression was performed over increasing lengths of time and the corresponding R^2^ value was plotted. A time window of five hours (red line) was selected as a systematic compromise for all conditions.

